# Towards Equitable MHC Binding Predictions: Computational Strategies to Assess and Reduce Data Bias

**DOI:** 10.1101/2024.01.30.578103

**Authors:** Eric Glynn, Dario Ghersi, Mona Singh

## Abstract

Deep learning tools that predict peptide binding by major histocompatibility complex (MHC) proteins play an essential role in developing personalized cancer immunotherapies and vaccines. In order to ensure equitable health outcomes from their application, MHC binding prediction methods must work well across the vast landscape of MHC alleles. Here we show that there are alarming differences across individuals in different racial and ethnic groups in how much binding data are associated with their MHC alleles. We introduce a machine learning framework to assess the impact of this data disparity for predicting binding for any given MHC allele, and apply it to develop a state-of-the-art MHC binding prediction model that additionally provides per-allele performance estimates. We demonstrate that our MHC binding model successfully mitigates much of the data disparities observed across racial groups. To address remaining inequities, we devise an algorithmic strategy for targeted data collection. Our work lays the foundation for further development of equitable MHC binding models for use in personalized immunotherapies.

## Introduction

Immunotherapies, where individuals’ immune systems are harnessed to attack cancer cells, are amongst the most exciting new approaches for cancer treatment^1^. T cells, which are central components of this process, detect and then either eliminate cells that display foreign or mutated peptides in complex with MHC molecules on their surfaces or secrete signals to stimulate other immune cells. As MHC proteins are highly polymorphic across human populations and each allele can bind a different set of peptides^2^, machine learning methods to predict which peptides each MHC protein can bind^3^ have become essential in screening cancer proteomes for potential T cell epitopes^4,5^.

*In silico* T cell epitope screens utilize MHC-peptide binding predictions to identify cancer neoantigens, which are peptides generated from tumor-specific mutations that are presented by an individual’s MHC molecules. Landmark advances in cancer immunotherapy, brought about with checkpoint blockade inhibitors, have resulted in therapeutics that can overcome the multiple escape mechanisms that tumors can use to evade T cell-mediated elimination^5^. These advances have propelled neoantigen discovery and the development of personalized neoantigen vaccines–numerous of which are in clinical trials–to therapeutically target tumors with a level of specificity unrivaled by existing cancer treatments^7^. The improvement of computational methods for predicting MHC-peptide binding is frequently cited as one of the greatest limitations, and areas of need, for the efficacy of emergent neoantigen immunotherapies^8^.

A major challenge in predicting MHC-peptide binding is that while there are over 13,000 distinct class I MHC protein allelic variants^9^, binding data is only available for a relatively small number of MHC alleles. Pan-MHC models leverage the similarity between MHC alleles and their binding repertoires to make binding predictions for any MHC allele^10-16^. However, a major limitation of current approaches is that predictive performance can only be estimated for the small minority of MHC alleles with sufficiently large binding datasets. Since MHC binding algorithms influence the efficacy of downstream biotechnologies^8,17,18^, it is absolutely necessary to understand the quality of MHC-peptide predictions for any allele. Moreover, because the frequency of MHC alleles varies across different human subpopulations^19^, there is substantial danger that differing MHC-peptide prediction performance across alleles may yield disparate outcomes in immunotherapies across racial and ethnic groups, thereby exacerbating the already significant disparities in cancer mortality and treatment across racial groups^20^. Existing approaches to predict MHC-peptide binding^21^ have not been designed or comprehensively evaluated with respect to equitable performance across individuals and populations.

Here we introduce a framework to enable the development of equitable class I MHC binding prediction models. We begin by highlighting the importance of developing MHC binding models with the explicit goal of equitable predictions by demonstrating that there are stark disparities in how much binding data is associated with the MHC alleles of different individuals and of individuals across racial and ethnic groups. We next introduce a framework for equitable MHC models based upon the co-development of a deep learning model MHCGlobe for predicting peptide-MHC binding, along with a novel machine learning method MHCPerf to estimate how well this pan-MHC model performs on unseen MHC alleles. We demonstrate that MHCGlobe has state-of-the-art-performance in predicting peptide-MHC binding as well as high estimated performance for the majority of MHC alleles, despite most of them having little or no binding data. These performance estimates across all alleles–even those with no existing binding data– comprise, to the best of our knowledge, the first analysis of how well pan-MHC models support the diversity of MHC alleles found across human populations. Despite excellent overall per-allele performance, we find that MHCGlobe nevertheless exhibits some performance differences across MHC alleles and this leads to disparate performance estimates when applied to individuals from different racial groups. Finally, we devise algorithms to select MHC alleles to prioritize for future data collection as a strategy to address the effect of data imbalances on MHC binding model performance.

The current study contributes multiple resources to clinicians, immunologists, and computational researchers whose work intersects with the adaptive immune response to MHC-bound antigens. For personalized vaccine design, our allele-level performance estimates can be used to prioritize neoantigens for which predictions are expected to be of higher quality. For experimentalists, we prioritize MHC alleles for which data collection is estimated to have great utility in improving equitable performance across the landscape of underserved alleles. For researchers actively working to improve performance of machine learning models for MHC binding, we release the most comprehensive assessment of how the performance of a pan-MHC model is affected by data collection, including allele-level performance estimates revealing model “blind spots” that need to be addressed. Further, ethical questions pertaining to individual and group fairness are raised here and invite further work to establish metrics of fairness in the field of T cell antigen discovery and personalized therapies. Individually and combined, we hope these resources inspire future efforts to measure and mitigate inequality in MHC binding models and in T cell epitope discovery as well as to promote innovations to ensure both effective and equitable personalized therapies.

## Results

### Amount of binding data across human MHC class I alleles varies significantly across individuals and racial and ethnic groups

In order to address how well pan-MHC binding models may be expected to work across the landscape of documented human MHC class I alleles, we first assess the binding data available for pan-MHC models to learn MHC binding patterns from. Of the greater than 13,000 documented human MHC class I protein allelic variants (HLA-A, HLA-B and HLA-C genes), only 180 have any binding data (with 940,238 positive binding instances, and 1,119,627 total instances across all alleles [See Methods]). The number of positive binding instances ranges from one to over 160,000 per allele (Figure 1A). Pan-MHC models commonly represent each MHC allele with a pseudosequence comprised of 34 peptide-interacting residues within the MHC’s peptide binding pocket, yet only a small fraction of the distinct set of pseudosequences are associated with any binding data, corresponding to 5.56%, 5.14% and 2.41% of HLA-A, HLA-B and HLA-C pseudosequences, respectively (Figure 1B).

**Figure 1.**
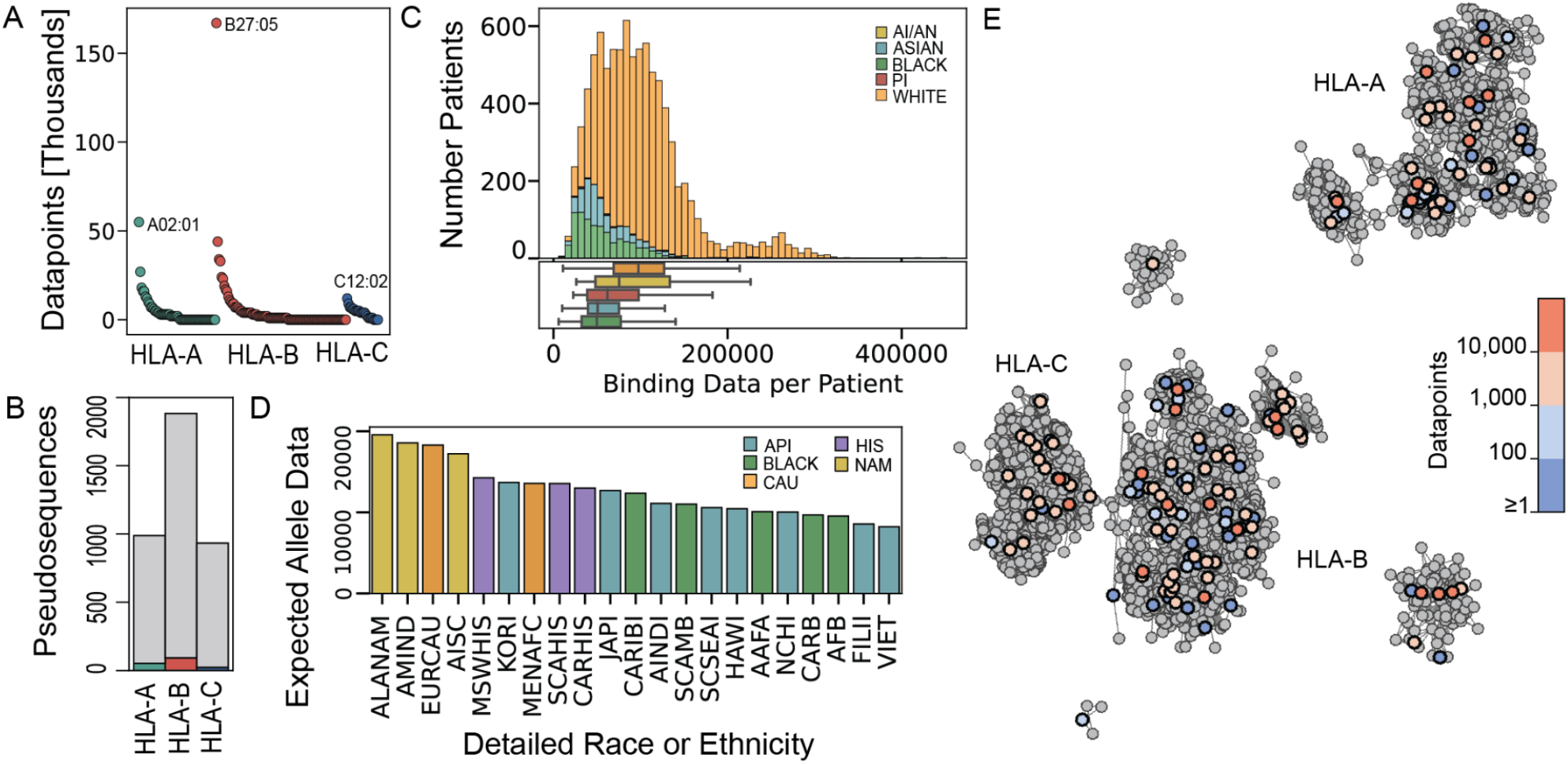
Variation in peptide-MHC binding data per HLA allele and across racial and ethnic groups. **(A)** The number of positive peptide-MHC binding datapoints for each of the 180 canonical HLA alleles with any data; each dot represents one allele. The allele with the most data for a given gene is labeled. **(B)** The number of distinct HLA pseudosequences with binding data (colored) or without (grey) per HLA gene. **(C)** (Top) A histogram of the total number of datapoints associated with the class I HLAs of each of the individuals from the TCGA, computed as the sum of the number of binding peptides known for each of their six class I HLA alleles. Colors represent the race categorization recorded for each patient by TCGA (AN/NA is “American Indian or Alaska Native”, Black is “Black or African American”, and PI is “Native Hawaiian or Other Pacific Islander”). (Bottom) Boxplots of the total amount of data per patient, separated by race. **(D)** For each of21 racial groups (see Supplementary Table 1 for descriptions), we give the expected number of positive binding instances for an allele sampled from that population according to the frequency with which that allele occurs in the population. Colors represent the broad race and ethnic categoriza tion associated with each of the detailed race or ethnic categorization (Native American (NAM) in gold, Caucasian (CAU) in orange, Hispanic (HIS) in purple, Asian and Pacific Islander (API) in blue, and Black in green). **(E)** Network visualization of 3,804 distinct MHC pseudosequences, where a node represents an individual MHC pseudosequence, and edges connect similar pseudosequences (see Methods). Pseudosequences without associated binding data are grey, and those with data are colored according to how much data are associated with them.

We next quantify the number of positive binding instances associated with the MHC class I alleles of patients from the The Cancer Genome Atlas (TCGA) project. We find highly variable coverage across individuals profiled in TCGA—ranging from 5,783 to 452,277 binding instances per patient, with a median of approximately 89,000 instances (Figure 1C). By recorded race, White individuals tend to have significantly more data associated with their MHC alleles than do Asian or Black individuals (*p*-values of 1.15e-141 and 3.93e-196, respectively, Mann-Whitney test). To extend this analysis, we next utilize MHC allele frequencies for 21 racial and ethnic populations (estimated from HLA typing of 2.90 million individuals^19^) to calculate the expected amount of binding data for an allele sampled from each population. Corroborating the pattern observed in TCGA patient coverage, Caucasian and Native American populations are expected to have more data associated with a randomly sampled MHC allele as compared to Hispanic, Asian, and African American populations (Figure 1D). As pan-MHC machine learning models share data across alleles in order to make predictions for alleles with limited or no binding data, we next visualize data coverage across alleles in the context of their relationships to one another using a network (Figure 1E) where MHC alleles that are sequence-similar to each other are connected by an edge (see Methods). We find that 94.3%, 99.4% and 99.7% of MHC alleles are, respectively, adjacent to, one hop away from and two hops away from a MHC allele with data; this suggests that pan-MHC models should be able to leverage data across alleles to make good predictions for all alleles.

### Novel framework assesses allele-level performance of pan-MHC models

Having shown that there vast differences in the amount of binding data associated with each MHC allele, we next introduce a framework that both develops a pan-MHC model as well as uncovers how allele-level performance for diverse MHC alleles is affected by the availability of binding data. In particular, we devise two systems: (1) MHCGlobe, a state-of-the-art deep learning method for pan-MHC binding affinity prediction; and (2) MHCPerf, a novel method for estimating allele-level performance of MHCGlobe (Figure 2). MHCGlobe takes as input a query MHC class I allele’s pseudosequence and a peptide sequence (8-15 amino acids in length) and outputs a score predicting the MHC-peptide binding affinity value (Figure 2A). MHCGlobe is an ensemble of deep neural networks trained on MHC-peptide binding data. MHCPerf (Figure 2B) is a neural network model that takes as input a query MHC allele’s pseudosequence as well as MHCGlobe’s training data to predict the Positive Predictive Value (PPV), which we use to measure the expected performance MHCGlobe would achieve on a test binding dataset for the query MHC allele (See Methods). For a query MHC allele, PPV estimates the fraction of the top peptides predicted by MHCGlobe to bind it that actually do; we use PPV to estimate per-allele performance as in most applications of MHC binding models, the top predictions are considered for downstream analysis and only actual binding pairs are of interest. MHCPerf considers the relationship of the input query MHC pseudosequence to the MHC alleles used in training MHCGlobe, along with the amount of data associated with each allele in the training set. In order to train MHCPerf, we simulate the data collection process and measure changes to allele-level performance as a result of training MHCGlobe on numerous incrementally updated datasets (See Methods). In principle, an approach such as MHCPerf can be used to predict the performance of any existing pan-MHC model, as long as it is computationally feasible to train the pan-MHC model many times. Together, MHCGlobe and MHCPerf provide a level of transparency unavailable for existing MHC binding prediction methods and will allow researchers to anticipate when model performance may be a limitation for specific MHC alleles of interest, and when variation in performance across alleles may confound downstream bioinformatic analyses.

**Figure 2.**
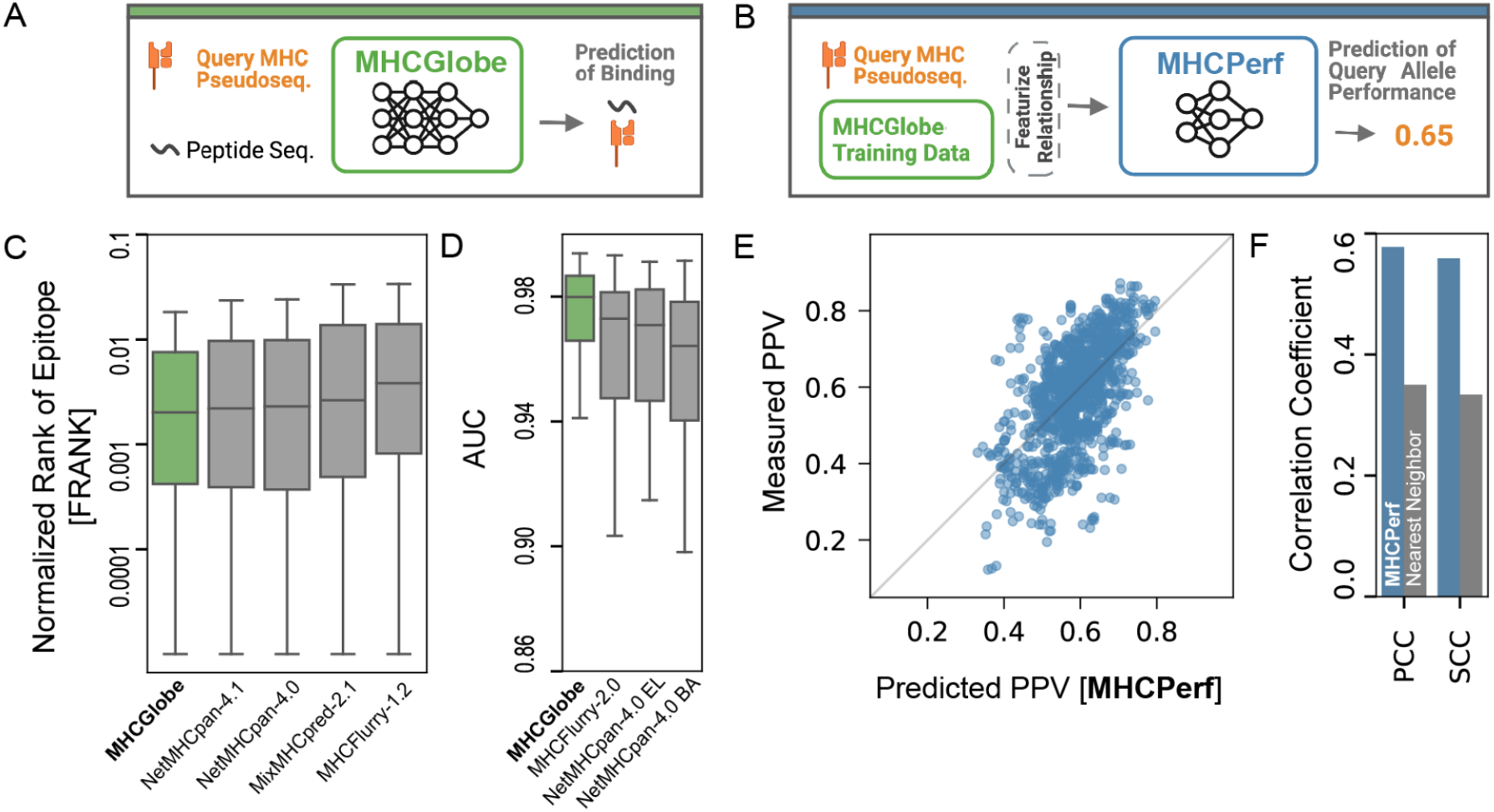
Schematic of new methods and performance summary. **(A)** MHCGlobe is a pan-MHC model that predicts the binding affinity value for an input MHC-peptide pair. **(B)** MHCPerf predicts an expected performance value (using PPV as the performance metric) that MHCGlobe would achieve for an input query MHC allele and specific MHCGlobe training dataset (see Methods). **(C)** Comparison of MHCGlobe to other pan-MHC methods when evaluated on the FRANK benchmark dataset of documented CD8+ T-cell epitopes published with NetMHCpan 4.1 (Reynisson 2020). Lower FRANK values indicate that the documented epitope was ranked as a stronger MHC binder compared to other peptides from the same source protein (See Methods). MHCGlobe has the lower median FRANK scores for the correct epitope than do the other methods, and it has significant improvement over MixMHCpred and MHCFlurry (one-sided Mann-Whitney p-values l.3e-05 and 8.Se-16, respectively). **(D)** Comparison of MHCGlobe to other pan-MHC methods when predicting MHC-peptide binding for monoallelic cell lines (published with MHCFlurry 2.0 O ‘Donnell 2020), as measured by Area Under the Receiver Operator Curve (AUC). MHCGlobe performs significantly better than MHCFlurry-2.0 BA, NetMHCpan-4.0 EL and NetMHCpan-4.0 BA (one-sided Mann-Whitney p-values of2.6e-3, 9.6e-4, and 3.7e-6, respectively). **(E)** For each MHC allele unseen during its training (10-fold cross validation, all data for an allele in one fold), MHCPerf ‘s output PPV (x-axis) as compared to the actual measured PPV of MHCGlobe for that allele when using data from the other nine folds (y-axis). Grey diagonal line indicates where measured PPV equals predicted PPV. **(F)** The Spearman and Pearson correlation of MHCPerf ‘s predicted PPVs and MHCGlobe ‘s actual PPV (blue) as compared to the correlations obtained when predicting PPVs based on using tl1e actual PPV of tl1e test allele ‘s nearest training fold neighbor (grey).

### MHCGlobe has state-of-the-art performance, and MHCPerf accurately estimates per-allele performance

To ensure the usefulness of MHCGlobe and MHCPerf to the broad research community, we start by evaluating MHCGlobe’s performance against performance of the available alternative methods NetMHCpan-4.1^11^, NetMHCpan-4.0^12^, MixMHCPred-2.1^14^, MHCFlurry-1.2^15^ and MHCFlurry 2.0^16^. We evaluate MHCGlobe performance on two benchmarks published with NetMHCpan4.1 and MHCFlurry 2.0, enabling comparison to alternative methods that were benchmarked previously; results shown here for other approaches are as reported in these previous benchmarks (see Methods). We first assess how well each method performs in identifying 1,660 known T Cell (CD8+) epitopes from full protein sequences. Here, for each documented CD8+ T Cell epitope, we use each of the methods to rank the predicted binding affinity of the epitope amongst all other 8-14-mer peptides from the same source protein sequence. MHCGlobe outperforms all other methods, giving stronger binding affinities than the other methods to the known epitopes within the sequences (Figure 2C). Next, we consider how well each of the methods predict MHC-presented peptides for 100 monoallelic engineered cell lines. MHCGlobe outperforms all other binding affinity prediction methods in classifying peptides that either bind or do not bind the tested alleles (Figure 2D).

To evaluate our second model, MHCPerf, we use 10-fold cross-validation and test MHCPerf’s ability to predict PPV for HLA alleles that it has not seen in its training (Figure 2E). Predicted PPV by MHCPerf is highly correlated with measured PPV values across the allele-level performance measurements, with Pearson’s Correlation Coefficient (PCC) of 0.58 and Spearman’s Correlation Coefficient (SCC) of 0.56. To our knowledge, MHCPerf is the first machine learning method that predicts allele-level performance of a pan-MHC model. Thus, we are not able to compare MHCPerf to previous approaches; however, Nielsen *et. al*.^10^ showed that the measured performance of an allele is positively correlated with that of its most sequence similar MHC alleles with available training data. We use this observation to create a “nearest neighbor” approach which predicts the PPV of a query allele as the measured PPV of the most similar allele with training data. MHCPerf greatly outperforms the nearest neighbor approach (PCC of 0.58 versus 0.35 and SCC 0.56 versus 0.33 for MHCPerf versus Nearest Neighbor) (Figure 2F). Thus, MHCPerf newly offers the ability to reliably estimate allele-level performance for previously uncharacterized HLA alleles (i.e., those with no experimentally measured binding data), and opens up new opportunities to assess the quality of MHC-peptide predictions across diverse sets of individuals.

### MHCGlobe mitigates disparities across racial and ethnic populations but inequities remain

We next use MHCPerf to quantitatively assess MHCGlobe’s predicted performance across alleles, individuals, and racial and ethnic groups; to our knowledge, this is the first such assessment of a pan-MHC model. Remarkably, MHCGlobe has reasonably high estimated performance for the majority of alleles (median predicted PPV 0.65), which is notable considering that 87% of the MHC pseudosequences are not associated with any binding data. However, we observe large variation in MHCGlobe’s estimated performances, ranging from PPVs of 0.27 to 0.94, across the 3,804 distinct MHC pseudosequences (Figure 3A). Consistent with the distribution of the known binding data, pseudosequences corresponding to HLA-B alleles overall have the highest predicted performance, followed by HLA-A and then HLA-C alleles. For each individual profiled in TCGA, we can compute the estimated performance of all of their MHC alleles by summing the MHCPerf estimated PPVs across their six alleles. We see a clear spread in performance scores across individuals, ranging from 3.6 to 4.4 (Figure 3B). Notably, MHCGlobe’s pan-MHC model partly mitigates the data disparity, with the differences between individuals with White and Asian ancestries and between individuals with White and Black ancestries not as statistically significant when comparing MHCGlobe’s expected performance as they are when comparing the amount of available data (compare with Figure 1C). However, we still observe significant differences in the expected performances of MHCGlobe across populations, and this overall pattern is corroborated when computing the expected performance estimate for a random HLA allele selected from each of the 21 racial and ethnic groups assessed earlier for data coverage (Figure 3C). Based on these assessments, the effect of data sharing in pan-MHC modeling is unlikely to completely mitigate existing data coverage inequalities across individuals, races and ethnic groups—and indicates that efforts towards fair pan-MHC binding predictions methods and downstream technologies should prioritize strategic data collection to address these disparities.

**Figure 3.**
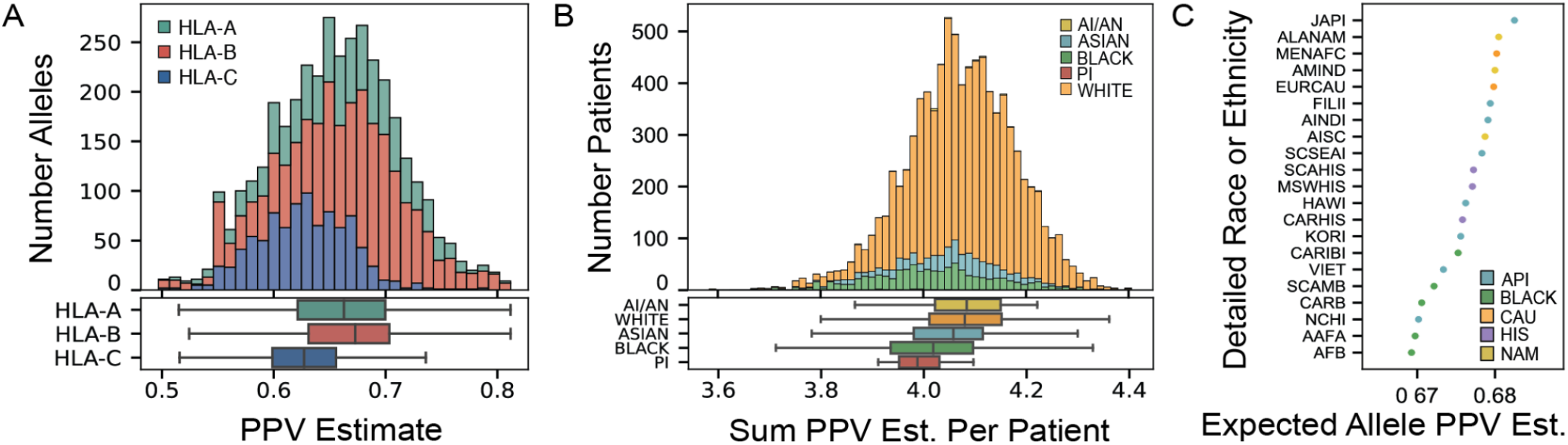
Assessment of MHCGlobe ‘s predicted performance per-allele, per-individual, and per-broad racial group. **(A)** For each of 3,804 distinct MHC pseudosequences, MHCPerf predicts the expected PPV that MHCGlobe will have on a test set of data for that allele. Shown is a histogram (top) and boxplot (bottom) of the predicted PPVs. Expected performance differs between HLA genes (Mann-Whitney *p*-values of 0.003, 2.3e-41, 6.7e-76 for comparisons between HLA-A and HLA-B, HLA-A and HLA-C, and HLA-B and HLA-C, respectively). 72 alleles with outlier values defined as ± 1.5 times the interquartile range are not shown. **(B)** (Top) The number of individuals from The Cancer Genome Atlas (TCGA) with a given patient-level PPV score, which is computed as the sum of the PPV estimates corresponding to each of their six class-I HLA alleles. Colors represent the race categorization recorded for each patient. (Bottom) The distribution of patient-level PPV scores aggregated by race. There are significant differences in the expected performances of MHCGlobe, when comparing White and Asian populations, White and Black populations and White and Pacific Island populations (two-sided Mann-Whitney p-values of l.56e-12, 3.63e-50, and 0.004, respectively). **(C)** The expected PPV estimate for MHCGlobe, as estimated by MHCPerf, for an allele sampled from each of the 21 detailed racial and ethnic populations. Colors represent the broad race and ethnic categorization associated with each of the detailed race or ethnic categorizations as reported by Gragert et. al. (2013), as in Figure lD.

### An algorithmic strategy based on MHCPerf identifies HLA alleles for data collection in order to mitigate performance disparities across classical HLA alleles

To provide steps forward, we next utilize the capabilities of MHCPerf to (1) identify alleles with poor estimated performance in pan-MHC modeling and (2) identify key MHC alleles—dubbed MVP-MHCs (“Most Valuable Player” MHCs)—to prioritize for data collection in order to strategically improve performance broadly across numerous alleles and reduce inequities in pan-MHC models. In this section, all mentions of performance refer to estimated performance by MHCPerf. Visualizing the network of HLA pseudosequences colored by allele-level performance, rather than data, we observe distinguishable regions of the network with either strong or relatively weaker performance (Figure 4A). Regions of poor performance define “blind spots” of MHCGlobe in MHC sequence space which could be strategically addressed with collection of binding data for alleles in those regions.

**Figure 4.**
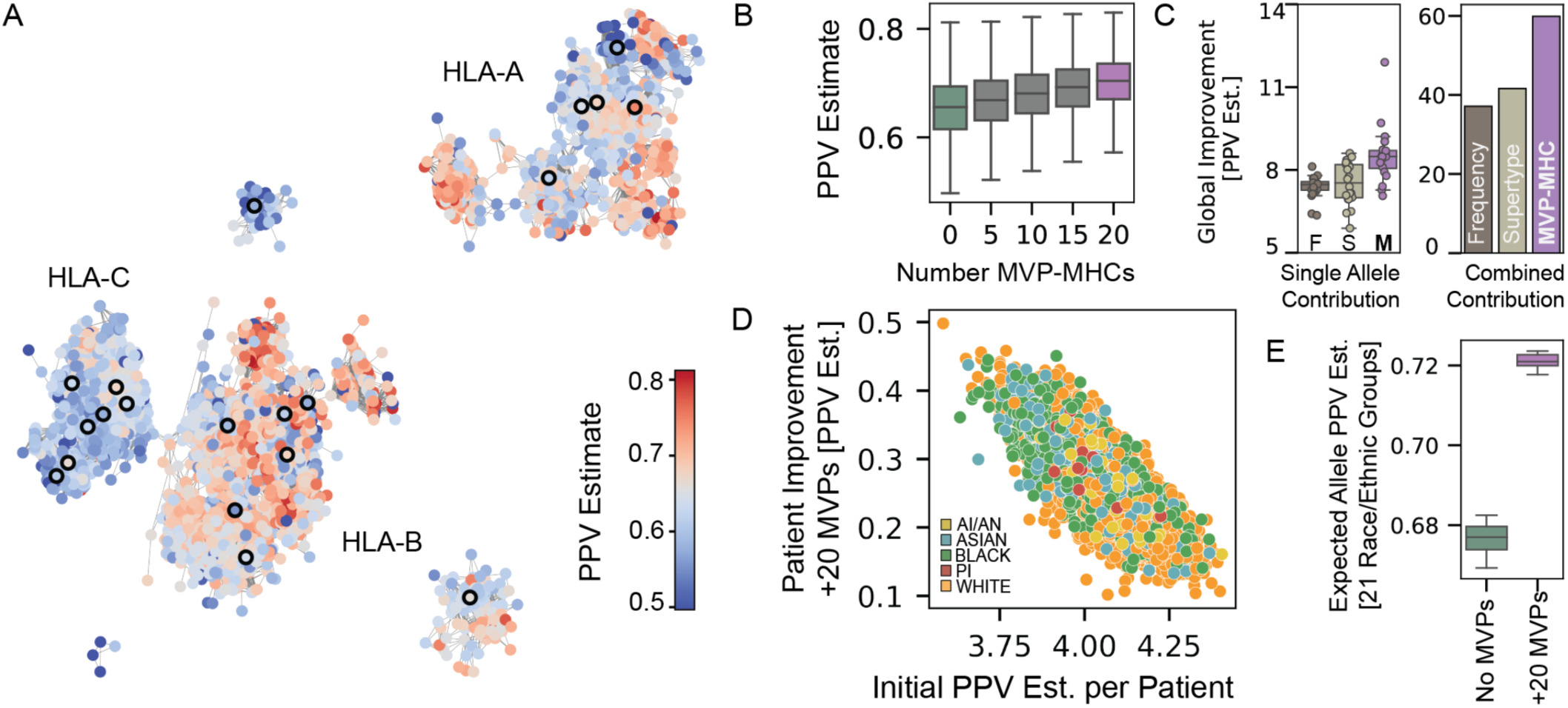
Towards fairness in pan-MHC modeling. **(A)** Network of distinct HLA pseudosequences and their relationships as in Fig 1C, but colored by their estimated PPV. Nodes with a thick black border represent the pseudosequences of first 20 MVP-l\1HC alleles selected for data prioritization. Color scale is limited to +/-1.5 times the interquartile range (IQR) to accentu ate contrast among the majority of alleles, meaning outlier values are colored the same as those with values at the boundary of +/-1.5 * IQR. **(B)** Incremental addition of 4,000 data points for each of the top 5, 10, 15 or 20 MVP-l\1HCs, followed by retrain ing l\1HCGlobe, is expected to improve performance across HLAs. The purple boxplot on the right corresponds to adding data for all of the top 20 MVP-l\1HC alleles, and then retraining l\1HCGlobe. The green boxplot on the left corresponds to the scenario where no alleles are given additional data.**(C)** Comparison of three approaches to select MHC alleles to prioritize for data collec tion as assessed by predicted Global PPV Improvement for all distinct HLA pseudosequences. Global PPV Improvement is defined as the sum across all 3,804 distinct HLA pseudosequences of the difference between PPV estimates after versus before adding 4,000 hypothetical data points to the specified alleles. The three approaches to allele selection are Frequency (Dark Brown); Supertype (Grey), and MVP-l\1HC (Purple). (Left) Global improvement in PPV estimates with the hypothetical data given to a single selected MHC allele. For each of the three approaches for allele selection, boxplots give the predicted Global PPV Improvement when adding data for one of the 20 selected alleles. The MVP-l\1HC approach leads to better predicted Global PPV Improvement (Mann-Whitney p-values of 4.16e-5 and 4.6e-4 comparing MVP-MHC to Frequency or Supertype, respec tively). (Right) Predicted Global PPV Improvement if data were added for all 20 selected alleles by each corresponding approach. **(D)** Improvement in individual-level PPV score (as computed in Figure 3B for individuals profiled in TCGA) after adding 4,000 data points to all of the top 20 MVP-MHC alleles plotted against the current patient-level PPV-estimate without additional data for MVP-l\1HC alleles. Colors correspond to patient race (as in Fig. ID). Patients with lower initial PPV estimates (x-axis) tend to have larger improvements (y-axis) in their patient-level PPV score. **(E)** Comparison of expected allele PPV estimates for the 21 race and ethnic groups (from Figure ID) when 4,000 data points are added to the top 20 MVP-l\1HC alleles or not. Adding data for the MVP-l\1HC alleles increases expected performance for l\1HCGlobe as well as reduces the variation across race and ethnic groups.

We therefore develop a greedy approach that leverages MHCPerf to tackle the problem of identifying MVP-MHC alleles, which if additional data was collected for, would have the most “beneficial” impact in improving MHCGlobe “weak spots” in performance. The greedy algorithm iteratively selects the MVP-MHC alleles that best improve the performances for alleles where MHCGlobe is underperforming (see Methods). We use our algorithm to select the top 20 MVP-MHC alleles to prioritize for strategic data collection to demonstrate the advantage of our approach. Incremental addition of these selected 20 MVP-MHC alleles raises the distribution of PPV estimates across alleles (Figure 4B). Moreover, the spread in the estimated performances across alleles declines, with the standard deviation decreasing from 0.062 initially to 0.053 after data for 20 alleles is added.

We highlight the putative advantage of selecting MVP-MHC alleles chosen by our algorithm compared to two alternative strategies, population frequency-based or the supertype-based prioritization of alleles. The population frequency-based approach chooses the 20 most common HLA alleles for target data collection (as estimated across 497 global population samples^22^). The supertype-based approach selects 20 alleles for prioritization by choosing one allele from each of 20 HLA “supertypes” associated with the least amount MHC binding data; MHC alleles are all classified into supertype clusters based on their sequence similarity^23^. To compare each data collection strategy, we use MHCPerf to estimate how much performance would be improved globally (across the 3,804 HLA pseudosequences) if 4,000 additional binding records were collected per allele selected by these approaches (Figure 4C). The global improvement across the pseudosequences, computed as the difference in the sum of PPVs across alleles, is greater for MVP-MHC alleles, considered individually, as compared to alleles selected by either of the two alternative methods (Figure 4C, left panel). This improvement is amplified considering the impact of adding data for all 20 alleles together (Figure 4C, right panel). Remarkably, prioritizing the top 20 MVP-MHC alleles is expected to improve patient-level performance for all of the TCGA patients, but yields the greatest improvement for patients currently with poorest performance (Figure 4D). We next assess the impact of adding data for the MVP-MHC alleles on estimated performance across racial groups and find that the expected performance of MHCGlobe across the 21 racial and ethnic groups (see Methods) is higher with additional data for the MVP-MHCs (Figure 4E). Moreover, the variability narrows across groups, which suggests that obtaining data for these prioritized MVP-MHCs should yield more equitable predictions of MHC binding across racial groups.

## Discussion

Computational methods for predicting MHC binding play a critical role in personalized cancer immunotherapies and effective vaccine design. In order to ensure that cutting-edge treatments benefit individuals from diverse racial and ethnic groups, it is essential that MHC binding models perform equitably across populations. Here we demonstrate that peptide-MHC binding data is highly imbalanced across the human HLA alleles, and this results in disparate data coverage across individuals and across racial and ethnic populations. To assess the impact of this data imbalance, we paired the development of a pan-MHC method, MHCGlobe, for predicting MHC-peptide binding with a new approach, MHCPerf, that estimates MHCGlobe performance for all MHC alleles, including those with no binding data. We demonstrate state-of-the-art performance for MHCGlobe and show that MHCPerf is able to reliably predict allele-level MHCGlobe performance values for alleles unseen in MHCGlobe’s training, only requiring the pseudosequence for the query MHC allele along with the training data used to train MHCGlobe.

While MHCPerf uncovers that MHCGlobe has high estimated performance for the majority of alleles— even for those that are not associated with any binding data—it also finds considerable variation in MHCGlobe’s performances across alleles, which leads to inequity in the quality of MHC-peptide binding predictions across racial and ethnic groups. Since MHCGlobe is trained on similar data as previous approaches, we expect that all widely used MHC class I predictors available today are likely to perform better as a whole for individuals with White ancestry than they do for individuals with Black or Asian ancestry. In theory, a framework like MHCPerf can—and should—be utilized to estimate the per-allele performance of any machine learning method for MHC-peptide binding, as long as it is computationally feasible to train the method on numerous subsets of the training data and measure performance.

To mitigate inequity in pan-MHC model performance, we propose a heuristic strategy for targeted data collection; our analysis suggests that such a strategy would lead to more equitable predictions by improving predictions more for underperforming alleles. In the future, it may be fruitful to define specific objective functions (e.g., to improve performance for specific sets of alleles for which predictions are particularly poor, or to improve performance for certain populations) and develop general optimization approaches to find solutions. Overall, identifying MHC alleles for targeted data collection is an important approach for obtaining equitable MHC-peptide predictions.

Another parallel avenue to obtain more equitable MHC-peptide binding predictions is to try to modify the training of neural networks. Here, all training data contributed equally to the loss function used for training. It may be beneficial to modify training so as to upsample or upweight data for less well-studied MHC alleles^24,25^. However, since alleles also have varying amounts of sequence similarity to each other, and sequence-similar alleles tend to have relatively similar binding profiles, it is likely that more specialized schemes that consider the similarity of alleles to each other will be necessary. Alternatively, the loss function used here minimizes error across all data points; the use of other loss functions that prioritize different goals may also be beneficial for obtaining more equitable predictions^26^. For any modification in training, it will be necessary to first verify that the overall performance of the model is state-of-the-art and then estimate (using an approach such as MHCPerf) allele-level and population-level performance in order to characterize whether MHC predictions are equitable.

In conclusion, our work has highlighted the importance of intentionally developing pan-MHC models that not only have state-of-the-art performance but also work well across diverse sets of alleles. We have presented an initial approach to estimate per-allele performance and have shown how it can be used to assess equity in MHC binding predictions and guide targeted data collection. The approaches introduced here lay the groundwork for further development of equitable MHC binding models, a key step in ensuring that future advances in personalized immunotherapies benefit genetically diverse individuals.

## Methods

### Peptide-MHC Binding Data

Our dataset is derived from the Immune Epitope Database (IEDB) (http://www.iedb.org/downloader.php?file_name=doc/mhc_ligand_full.zip) and the MHCFlurry 2.0 dataset^16^. Our dataset consists of class I MHC alleles, with a linear peptide epitope and peptide lengths between 8 and 15 amino acids. MHC-peptide binding data is either associated with a quantitative binding affinity (BA), or is qualitative (binding or not) and comes from elution experiments (EL) from either multiallelic (MA) or single-allelic (SA) cell lines. Preprocessing of the data, assignment of peptides from MA data to specific MHC alleles, and removal of duplicates is described in detail in the Supplement. After preprocessing steps, we have 64,616 BA, 441,648 MA and 537,229 SA positive MHC-peptide examples (full description in Supplementary Table 2). All positive qualitative records are assigned a quantitative affinity value of 100nM and measurement inequality of “<”, which indicates that positive examples should be assigned a value less than 100nM by the neural network^24^.

Synthetic negatives are generated by sampling *k*-mer amino acid subsequences uniformly at random from protein sequences in UniProt (downloaded 10/2020). For each allele, synthetic negatives are added at five times the total number of positive data points for that allele, with an equal number of different lengths (8-15-mers) represented in the sampled synthetic negatives. Synthetic negatives are assigned a quantitative value between 20,000-50,000nM, sampled uniformly, and measurement inequality of “>”.

MHC proteins are represented as pseudosequences, using the 34 amino acids in positions 31, 33, 48, 69, 83, 86, 87, 90, 91, 93, 94, 97, 98, 100, 101, 104, 105, 108, 119, 121, 123, 138, 140, 142, 167, 171, 174, 176, 180, 182, 183, 187, 191, and 195, as previously described^11,16^. MHC pseudosequences containing an unknown residue, “X”, in their 34-length pseudosequence are excluded.

### Construction of HLA Pseudosequence Network

The network consists of 3,804 nodes representing distinct class I HLA allele pseudosequences (HLA-A, - B, -C only). Pseudosequences used in the current study were released with MHCFlurry 2.0^16^. Edges between nodes correspond to similar pseudosequences defined as having a BLOSUM62 distance within the range 0-0.1. The BLOSUM62 distance of two MHC pseudosequences A and B is defined as in Neilsen *et. al*.^14^ as 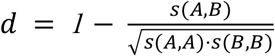 where *s*(*A, B*) is the BLOSUM62 alignment score^27^ of the pseudosequences. Omitted from the visualization are 11 of the 3,804 (0.3%) pseudosequences that are not similar enough to any other in our dataset to have a connecting edge in the network.

### Determining HLA haplotypes for TCGA data

Whole Exome Sequencing (WXS) data was downloaded for all normal (non-cancer) TCGA samples from the GDC Browser (https://portal.gdc.cancer.gov), and Optitype^28^ was used to determine HLA haplotypes. A preliminary processing step to Optitype is to use RazerS3^29^ to filter WXS reads to those able to align to the MHC allele reference library. Optitype was then run on each sample with default parameters. TCGA individuals with discrepancies across any of the six classical class I HLA alleles across multiple associated samples were excluded from this study. This resulted in high confidence HLA haplotypes for 8,942 individuals.

### MHCGlobe

#### Overview of MHCGlobe

MHCGlobe is an ensemble of neural networks, whose design and model selection is described below (see Ensemble Model Selection). Each neural network outputs a value between 0 and 1, corresponding to a log transformed binding affinity value (“nM” units), where measured affinity value, *a*, is transformed by 1-log(*a*)/log(50,000) as introduced by Nielson *et. al*.^30^. For training, we utilize a modified mean squared error loss that incorporates inequalities associated with each peptide-MHC pair so as to avoid penalizing predictions which fall within a specified range for binding affinity labels associated with semi-quantitative and qualitative binding records^16^. Each MHCGlobe neural network was trained using RMSprop gradient descent optimization within the TensorFlow (https://www.tensorflow.org/) training functions. In order to utilize binding affinity and elution data for peptide-MHC binding, a modified mean squared error loss function that handles measurement inequalities (“<”, “=”, ‘>”), introduced by MHCFlurry^15,16^, was used.

#### Peptide and MHC Input Representations

All peptides are represented as 15-mers in the manner introduced by MHCFlurry 1.2^15^. For peptides shorter than 15 residues, the first and last four residues are placed at the respective ends of the 15-mer and the central residues of the peptide are added to the center of the 15-mer representation. Unfilled positions in the 15-mer are set to an “X” residue. The representations of amino acids were treated as a hyperparameter, where each amino acid is represented either by a one-hot encoding (a vector of 20 zeros with a 1 in the position representing the amino acid), or by the corresponding row from BLOSUM62 which represents the similarity of that amino acid to all other amino acids. The unknown residue “X” is always encoded as a vector of 20 zeros.

#### Overview of Strategy for Choosing MHCGlobe Model Architecture

Our goal was to optimize MHCGlobe’s architecture without any exposure to the classical HLA genes, HLA-A, HLA-B, and HLA-C, which were reserved for the Leave-N-Out (LNO) cross validation (described in following sections, and which were used to measure how allele-level performance changes as a function of the binding data made available to MHCGlobe for training). Optimal hyperparameters were selected for the MHCGlobe ensemble by identifying neural networks trained on non-human MHC data that could best predict binding data for non-classical class I MHC alleles from human genes (i.e., HLA-E and HLA-G) (see Supplemental Figure 1). MHCGlobe’s architecture was thus fixed prior to exposure to any data from classical HLA alleles. In addition to having classical HLA data available for LNO cross validation, we reasoned that hyperparameters enabling a pan-MHC model to predict peptide-MHC binding for MHC alleles from unseen species and on non-classical class I MHC alleles would encourage the selection of hyperparameters that yield more generalizable models and thus are more desirable for a pan-MHC method.

#### Hyperparameter Optimization

Hyperparameter optimization was performed using Optuna^31^. Optuna optimizes hyperparameters by monitoring and minimizing mean squared error on a held out dataset, referred to here as the *optuna_set*. The space of hyperparameters is explored within an Optuna study, wherein Optuna iteratively samples and tests new hyperparameter combinations with sampling guided by performance measurements on the optuna_set from previous iterations. We used 100 iterations per Optuna study, and 15 Optuna studies in total. In total, this yielded 1500 explored hyperparameter settings from which we eventually chose individual deep neural network models to combine into an ensemble predictor. The hyperparameters we considered define neural network architectures and regularization parameters, and include the gradient descent algorithm’s parameters (see Supplemental Table 3).

For each Optuna study, the non-human MHC data was newly split into a training, validation, and optuna_set at a 4:1:1 ratio. The validation fold is used for early stopping when training the neural networks. Splits were created using distinct peptide-MHC pairs to avoid overlap between any two splits. Peptide-MHC pairs which are repeatedly observed in the data were always in the same data split. In order to ensure Optuna sufficiently tested neural networks of varying depths, we did not allow Optuna to sample the number of hidden layers as a hyperparameter for MHCGlobe; instead, the number of hidden layers was fixed to either 1, 2, or 3 layers and each tested for five of the 15 studies. All Optuna studies utilized a different splitting of non-human MHC binding data into training, validation, and optuna_set.

#### Ensemble Model Selection

To compose the ensemble, we iterated through each of the trained 1500 models and added the model to the ensemble which best improved ensemble performance on the non-classical human HLA data and repeated until no further improvement was achieved. The predicted binding score for an ensemble is taken as the arithmetic mean of each of the neural network predictions within the ensemble. Following this approach, we obtained an ensemble composed of three neural networks. Two of the selected neural networks have three fully connected hidden layers, and the third has two fully connected hidden layers. All three selected neural networks utilize at least one skip connection which allows information to flow through a given hidden layer and bypass the same hidden layer before being concatenated and passed to the following model layer. Since initialization values for weights and bias parameters of neural networks are important for learning and performance^32^, the initialization weights and biases for these three networks are set to be the same as they were prior to training on non-human MHC data and the same initialization weights and biases are used any time the particular neural network model is retrained. Once the ensemble model was selected, the full MHCGlobe model was trained on all available MHC binding data.

#### Early Stopping for MHCGlobe Training

For all MHCGlobe variants trained in this study, early stopping was used to prevent overfitting. MHCGlobe was trained for a maximum of 200 epochs, and halted by early stopping if performance on the validation set did not improve over 20 epochs by at least 0.0001. An allele-balanced training and validation set was created by partitioning 1/5th of the binding data for each allele to the validation set. As certain peptide-MHC records were observed multiple times in the available MHC binding data, training and validation partitions were performed using distinct peptide-MHC records, and repeated observations were added back into their respective partitions following partitioning.

### Benchmarking MHCGlobe Performance

We benchmarked MHCGlobe using datasets published with NetMHCpan 4.1^11^ and MHCFlurry 2.0^16^. In all benchmark comparisons to other methods, the MHCGlobe model was retrained after removing all peptide-MHC pairs that occur in the benchmark test set of interest. The NetMHCpan 4.1 benchmark^11^ consists of 1,660 protein sequences with documented CD8+ T-cell epitopes. For each documented CD8+ T Cell epitope, the epitope should have a higher binding affinity than other 8-10-mers peptides from the same source protein sequence. For each sequence, predicted binding scores of all peptides within it are rank normalized to obtain a FRANK score for each documented T cell epitope, where peptides with the highest predicted binding affinity are given a score of 0 and the peptides with the lowest predicted binding affinity are given a score of 1. Lower FRANK scores for the correct epitopes correspond to better performance. To make Figure 2A, FRANK scores for NetMHCpan 4.1, NetMHCpan 4.0, MixMHCpred 2.1, and MHCFlurry 1.2 were given in Supplementary Table 7 of paper^16^, and the FRANK scores for MHCGlobe were computed in this study using MHCGlobe binding predictions on the same benchmark. The MHCFlurry 2.0 benchmark^16^ consists of 100 monoallelic cell lines where peptides that either bind or do not bind the HLA allele are given^33^. For each sample, an area under the receiver operator curve (AUC) is computed to measure performance. To make Figure 2B, binding prediction scores for MHCFlurry 2.0, NetMHCpan 4.0 BA and EL were given in Supplemental Data S2 of paper^16^ and MHCGlobe binding scores were added to compute the AUC for MHCGlobe. While MixMHCPred was originally included in the benchmark of MHCFlurry 2.0 for a subset of MHC alleles, we exclude MixMHCPred here as we consider only pan-MHC methods that are able to make predictions for all class I MHC alleles in the benchmark.

#### Creating the MHCPerf Training Dataset via Leave-N-Out Cross Validation

##### Choosing MHCGlobe Training Data Subsets

A schematic of the LNO cross validation is given in Supplementary Figure 2. MHC alleles that had at least 50 distinct EL binding records (derived from either SA or MA data) were considered to have sufficient data to evaluate allele-level performance and were selected as test alleles. This yielded 125 test alleles. We generated 10 distinct MHCGlobe training sets for each test allele, where the first training set excluded data from the test allele. Successive training sets excluded all binding data from the next most similar allele to the test allele (using the BLOSUM62 distance defined above), such that at least 10 additional binding records had to be excluded; if fewer than 10 binding records were excluded, then an additional neighboring allele is excluded from the particular MHCGlobe training set. In total, 1,250 MHCGlobe training datasets were created.

##### Training Leave-N-Out Versions of MHCGlobe

A distinct version of MHCGlobe was trained for each of the 1,250 LNO training sets, following the same training protocol as the full model, including always being initialized with the same weights and bias values prior to being trained on binding data. The 1,250 models were trained in parallel using g3.4xlarge EC2 instances from Amazon Web Services (AWS).

##### Measuring Allele-Level Performance of MHCGlobe

For each of the 125 test alleles, MHCGlobe’s Positive Predictive Value (PPV) for that allele was measured for each of the 10 associated LNO MHCGlobe models. Each of the 125 test sets is composed of positive elution records of the corresponding test allele with the addition of synthetic negative peptides sampled at 99:1 with peptides of lengths 8-10 amino acids equally represented. PPV is computed as the proportion of positive labeled peptides in the top 1% of binding affinity predictions made by the corresponding LNO version of MHCGlobe.

#### Modeling Allele-Level PPV via MHCPerf

##### Featurization

The relationship between a query MHC allele and the binding data used to train MHCGlobe is used to create 63 features for MHCPerf that can be conceptually grouped into five categories (Supplementary Figure 2 and Supplemental Table 4). The first quantifies the BLOSUM62 sequence distance between the query MHC and each of the 10 most similar alleles with binding data included in the MHCGlobe training set. The second feature set quantifies the number of positive binding data points each of those 10 most similar alleles has in the training dataset. The third feature set quantifies the amount of positive binding data corresponding to training set alleles in one of eight sequence distance bins (0-0.1, 0.1-0.2, 0.2-0.3, 0.3-0.4, 04-0.5, 0.5-0.6, 0.6-0.7, 0.7+) representing the sequence distance between the query allele and the training set alleles. The fourth feature set quantifies the number of positive training data points corresponding to each of the 34 residue-position pairs of the query MHC pseudosequence. The fifth feature set is a single value quantifying the total number of positive binding records in the training dataset.

##### MHCPerf PPV-Balanced Folds

MHCPerf training data was split into PPV-balanced folds during the hyperparameter selection procedure and during 10-fold cross validation for MHCPerf performance evaluation. That is, because it is rare that MHCGlobe’s performance was either high or very low, in order to make each fold more representative of the full MHCPerf training set, we attempted to balance measured PPV across fold splits while keeping all instances corresponding to a given test MHC within the same fold. To do this, for each test allele, the mean PPV for MHCGlobe was first computed across the ten instances of LNO measurements. Then, test alleles were sorted by their mean LNO PPV and iteratively assigned to one of the folds.

##### MHCPerf Hyperparameter Selection

For MHCPerf, we utilized a single neural network. Hyperparameter selection was performed using the grid-search algorithm and a nested 3-fold cross-validation on the training dataset to select the best hyperparameters. Hyperparameters were chosen that gave the best average performance across the three folds. The neural network using the selected hyperparameters was then trained on the full training set (i.e., 9 folds), and used to predict MHCPerf performance for the alleles in the 10th fold (Supplemental Figure 2).

##### MHCPerf Trainings

MHCPerf was trained for 1,000 epochs. Minibatch training was employed with 10 records per batch. Early stopping was used and halted training if performance on the validation set did not improve over 100 epochs. The training and validation folds were generated with a random split, utilizing *validation_split=0*.*2* within the TensorFlow training function.

#### MVP-MHC Algorithm

The goal of the MVP-MHC algorithm is to identify key MHC alleles, for which collecting additional data would increase the expected performance of MHCGlobe on alleles for which it is currently underperforming. As MHCPerf and MHCGlobe make predictions on pseudosequences, the MVP-MHC algorithm is run on pseudosequences; for simplicity, we refer to each pseudosequence as an allele and assume that data would be collected for a single allele for a pseudosequence. The MVP-MHC (Most Valuable Player-MHC) algorithm utilizes MHCPerf to compute, for every allele, the effect of adding data for that allele on the performance of every other classical HLA allele. For each allele *j*, we compute an “impact” vector *vj* of dimensionality equal to the number of alleles, where the *i*-th element of the vector is equal to 0 if the estimated PPV of *i*-th allele is greater than the median estimated PPV across all alleles, and is otherwise set equal to the minimum of (1) the difference between the predicted PPV for allele *i* after adding 4,000 data points to allele *j* and the predicted PPV for allele *i* with the current data and (2) the difference between the median PPV for all alleles and the predicted PPV for allele *i* with the current data. In other words, this value is set to the amount of improvement for allele *i* when adding data for allele *j*, capped by how far the current performance for allele *i* is from the median performance. The total impact score for allele *j* is the sum of the elements of its impact vector. Intuitively, the impact vector for allele *j* measures how much closer the expected performance for each “underperforming” allele gets to the median

PPV with the addition of data for allele *j*; alleles that already have performance above the median performance are not considered when selecting an MVP. The MVP-MHC algorithm greedily picks MVP alleles. That is, at each iteration, the allele *m* with highest impact score is chosen as an MVP. Then for every other allele *j*, in order to choose MVP alleles that are complementary to each other, its impact vector is updated to be *v*_*j*_ - *v*_*m*_, with any negative entry of *v*_*j*_ subsequently set equal to 0. In the next iteration, these updated impact vectors are used to choose the next MVP-MHC. This MVP-MHC selection process is repeated until the desired number of MVP-MHC alleles have been selected or until all MVP-MHC candidates have been selected.

#### Data or PPV Estimate Expected Values for 21 Racial and Ethnic Groups

The expected data coverage (respectively PPV estimate) for the 21 detailed racial and ethnic groups shown in figures 1D and 3C is computed as *x*_*g*_ = ∑_*m*_ *c*_*m*_ · *f*_*g,m*_ where *x*_*g*_ the expected value of the amount of data (respectively PPV) for a given racial or ethnic group *g, c*_*m*_ is the amount of data (respectively PPV) for allele *m*, and *f*_*g,m*_ is the frequency of allele *m* in the given group (obtained from Gragert *et. al*.^19^. Allele frequencies within each group were normalized so that alleles for each of the associated HLA-A, HLA-B, and HLA-C genes together sum to one. Thus, the expected value represents the amount of data or PPV estimation for the average allele sampled from a given racial or ethnic population.

## Code Availability

Our predictors are freely available and open source (https://github.com/ejglynn/mhcglobe).

## Supplement

**Supplementary Figure 1.**
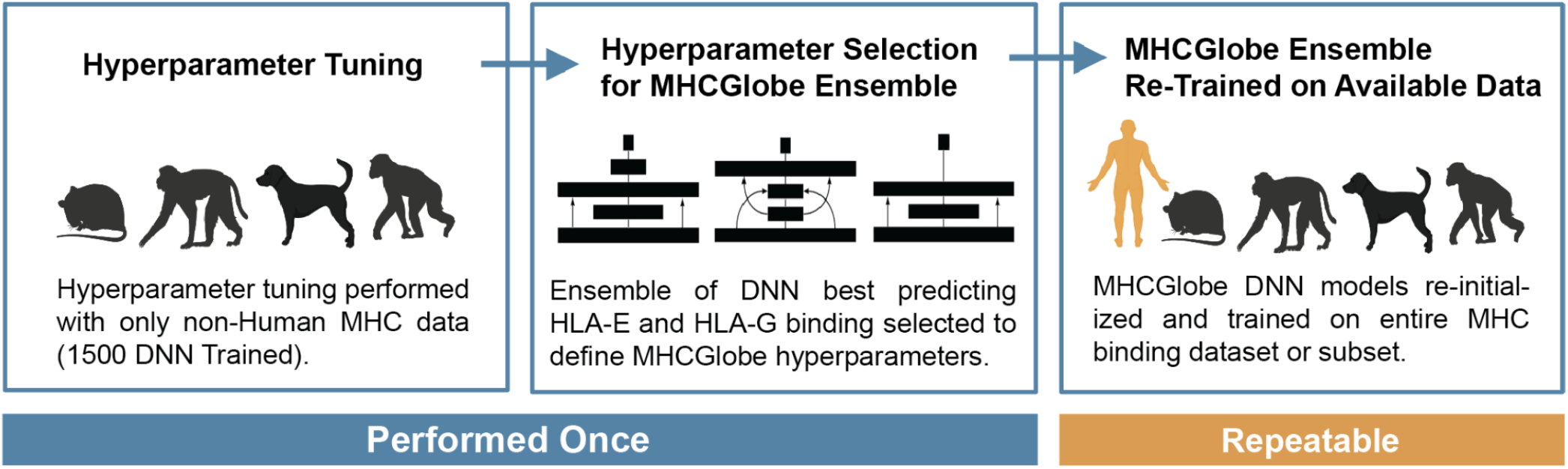
Schematic of novel strategy for pan-MHC model hyperparameter selection, ensemble construction, and training used for MHCGlobe. In order to enable the Leave-N-Out cross validation with MHCGlobe on classical class I HLA data, MHCGlobe hyperparameters were first optimized utilizing non-Human MHC binding data where 1500 distinct Deep Neural Networks (DNN) were trained. To assemble the best subset of DNN for the MHCGlobe ensemble model, DNN trained on non-Human data were added to the MHCGlobe ensemble based on performance of HLA-E and HLA-G binding data (non-classical HLA alleles). The DNNs selected for the MHCGlobe ensemble were fixed, enabling MHCGlobe to be re-trained with any subset of classical class I HLA alleles in the Leave-N-Out cross validation.

**Supplementary Figure 2.**
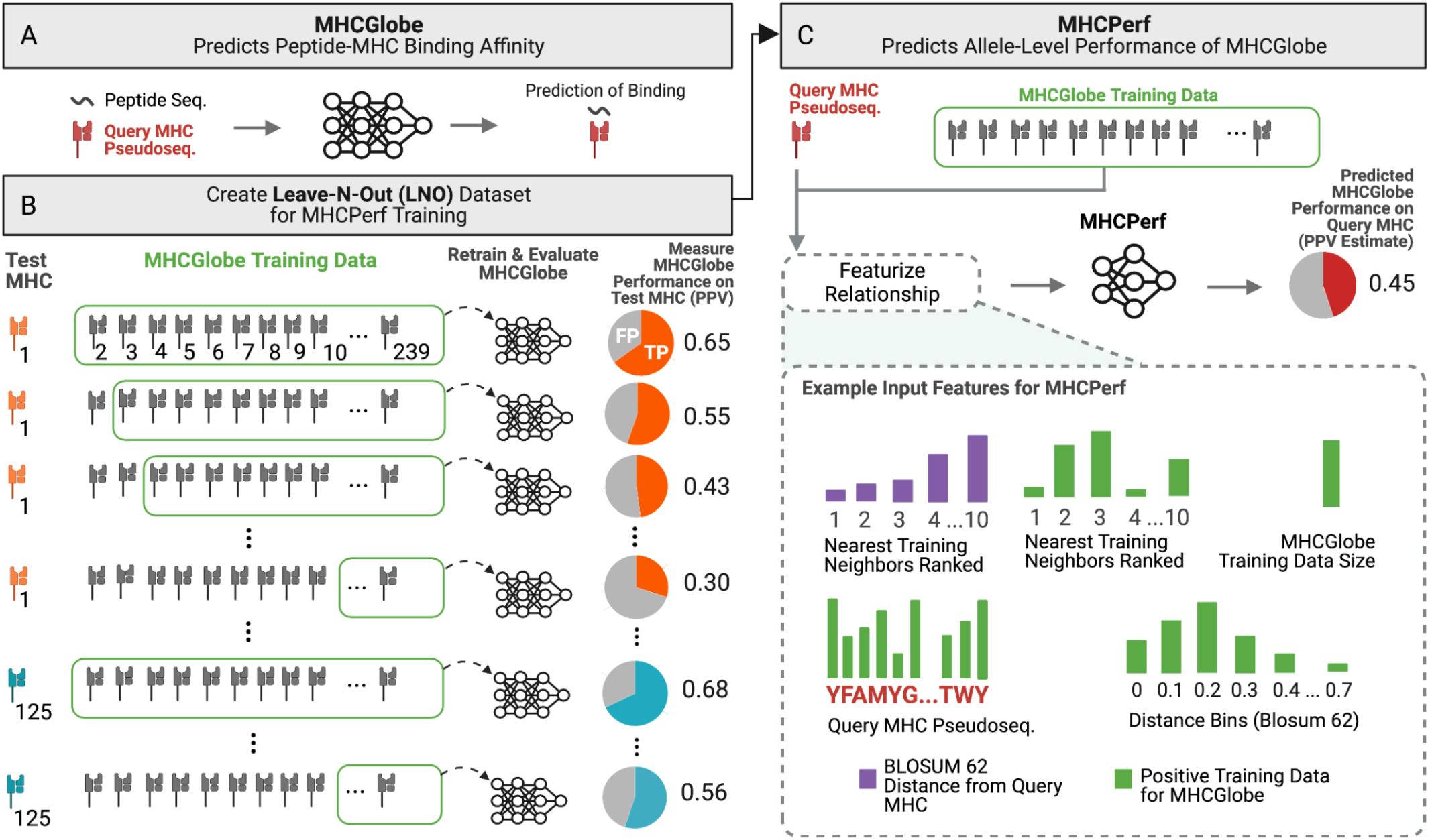
Schematic of new methods, creation of MHCPerf training dataset, and MHCPerf input features. (**A**) As in Figure 2, MHCGlobe takes as input a query class I MHC allele’s pseudosequence and a peptide sequence (8-15 amino acids in length) and outputs a score predicting the MHC-peptide binding affinity value. MHCGlobe is an ensemble of neural networks trained on MHC-peptide binding data. (**B)** Leave-N-Out cross validation to build a training dataset for MHCPerf. Each of 125 HLA alleles with sufficient data serves as a test MHC dataset for 10 rounds of cross-validation testing. MHCGlobe is trained on successively dissimilar MHC data by exclusion of neighboring alleles from the training dataset. For each of the 10 rounds, the actual performance of MHCGlobe on data for the test MHC is recorded. (**C**) As in Figure 2, MHCPerf takes as input a query MHC allele’s pseudosequence and uses MHCGlobe’s training data to predict MHCGlobe’s Positive Predictive Value (PPV) for that allele; this indicates the expected performance MHCGlobe would achieve on a test binding dataset for the query MHC allele (See Methods). (**D**) Schematic representation of MHCPerf’s feature sets used to describe the relationship between a query MHC pseudosequence and the MHCGlobe training data (see Supplemental Table 4).

**Supplementary Table 1.**
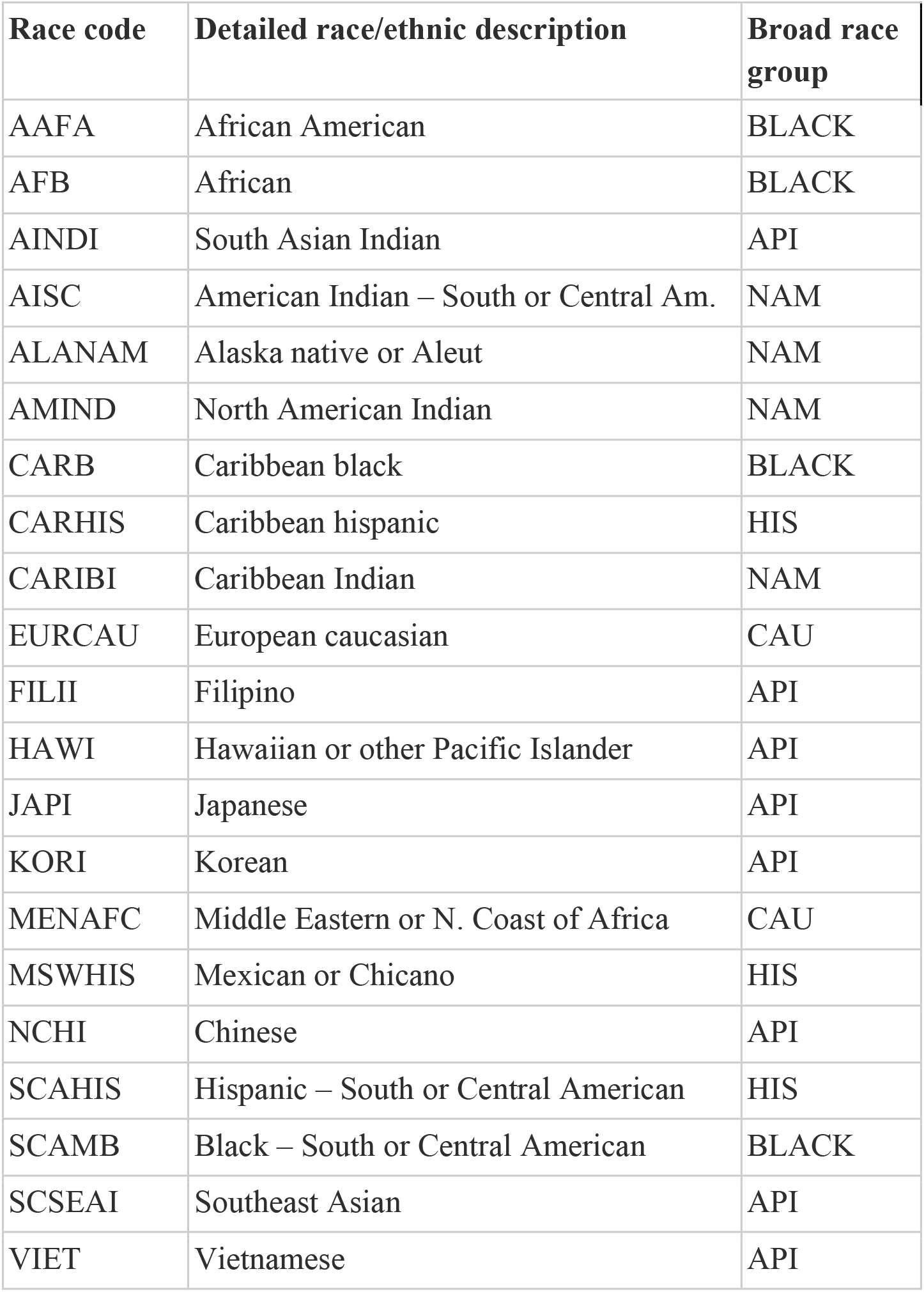
Labeling for 21 detailed racial and ethnic groups and corresponding to broad racial and ethnic groups for the allele frequency estimation published by Gragert *et. al*.^19^.

**Supplementary Table 2.** Number of Positive and Negative peptide-MHC records for Binding Affinity (BA), Single Allelic (SA) Elution, and Multi-Allelic (MA) Elution data.

|  | Positive | Negative |
| --- | --- | --- |
| BA | 64,616 | 138,646 |
| SA | 537,229 | 32,512 |
| MA | 441,648 | 37,576 |

**Supplementary Table 3:** MHCGlobe hyperparameters for each neural network in the ensemble model. Parameters listed from top to bottom correspond to: sequence encoding strategy for the peptide and the MHC pseudosequence; L1 regularization lambda value; the number of nodes for hidden layers 1, 2, and 3; boolean to use skip connection to add bypass connection over hidden layer 1 and/or hidden layer 2; epsilon and learning rate parameters for Root Mean Squared Propagation (RMSprop) gradient descent optimizer.

|  | Neural<br>Network 1 | Neural<br>Network 2 | Neural<br>Network 3 |
| --- | --- | --- | --- |
| **sequence_encoding** | ONE_HOT | ONE_HOT | ONE_HOT |
| **L1_reg** | 1 | 0 | 0 |
| **dense_units_1** | 128 | 256 | 256 |
| **dense_units_2** | 128 | 512 | 512 |
| **dense_units_3** | 512 | 128 | - |
| **skip_connection_1** | TRUE | TRUE | TRUE |
| **skip_connection_2** | TRUE | FALSE | NA |
| **epsilon** | 6.85E-07 | 5.52E-07 | 3.17E-07 |
| **rms_learning_rate** | 0.00113393 | 0.00103829 | 0.00191475 |

**Supplementary Table 4.**
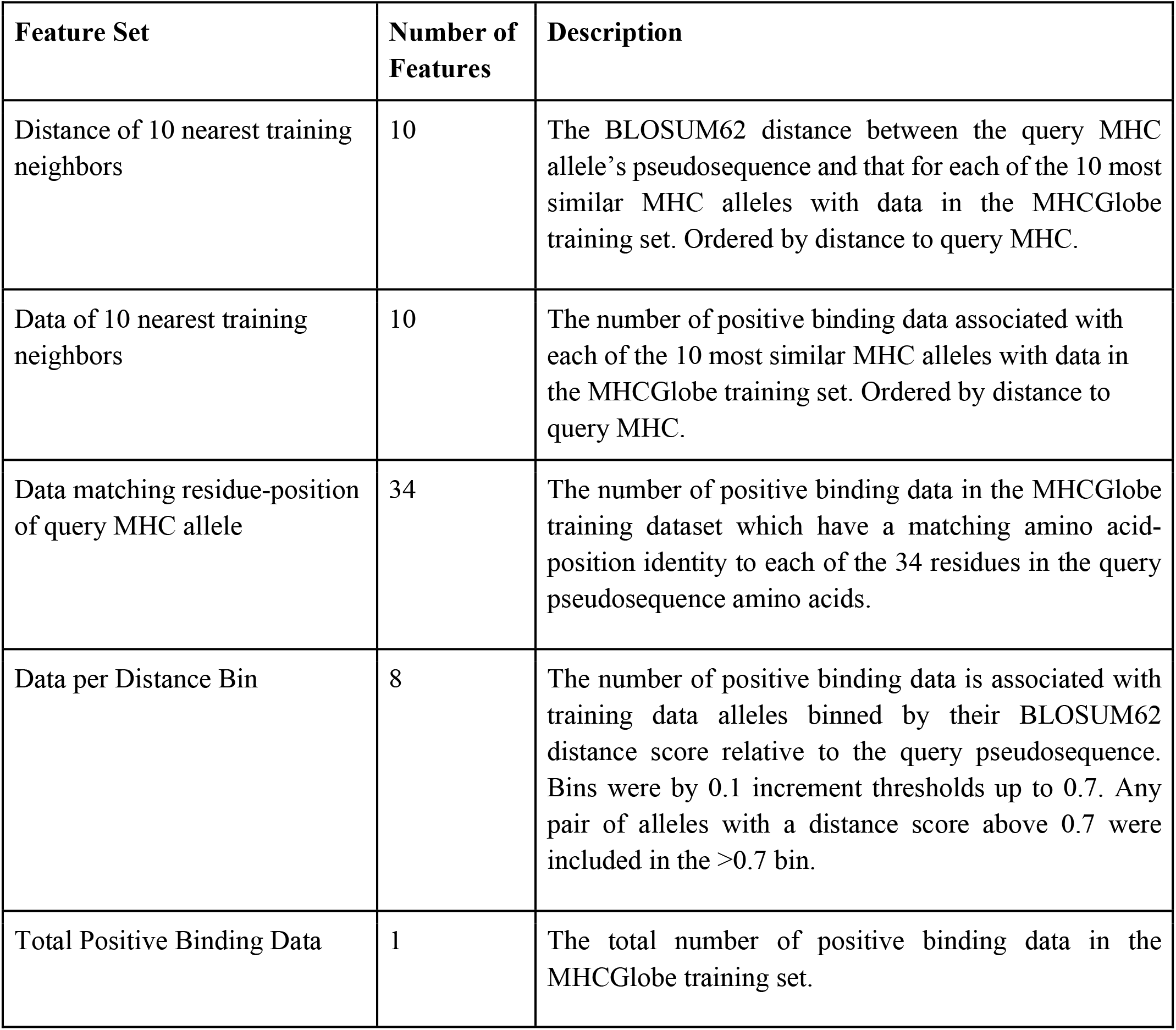
Descriptions of MHCPerf features. The input features of MHCPerf for a given query MHC were constructed by comparing the query MHC allele pseudosequence to different attributes of the MHCGlobe training set containing MHC binding data. For data counts only the number of positive binding data were utilized.

| Feature Set | Number of Features | Description |
| --- | --- | --- |
| Distance of 10 nearest training neighbors | 10 | The BLOSUM62 distance between the query MHC allele's pseudosequence and that for each of the 10 most similar MHC alleles with data in the MHCglobe training set. Ordered by distance to query MHC. |
| Data of 10 nearest training neighbors | 10 | The number of positive binding data associated with each of the 10 most similar MHC alleles with data in the MHCglobe training set. Ordered by distance to query MHC. |
| Data matching residue-position of query MHC allele | 34 | The number of positive binding data in the MHCglobe training dataset which have a matching amino acid-position identity to each of the 34 residues in the query pseudosequence amino acids. |
| Data per Distance Bin | 8 | The number of positive binding data is associated with training data alleles binned by their BLOSUM62 distance score relative to the query pseudosequence. Bins were by 0.1 increment thresholds up to 0.7. Any pair of alleles with a distance score above 0.7 were included in the >0.7 bin. |
| Total Positive Binding Data | 1 | The total number of positive binding data in the MHCglobe training set. |

## Preprocessing Peptide-MHC Binding Data

MHC epitopes curated by IEDB were downloaded at http://www.iedb.org/downloader.php?file_name=doc/mhc_ligand_full.zip. Instances were filtered to include only those with class I MHC alleles, with a linear peptide epitope and peptide length between 8 to 15 amino acids (i.e., *MHC allele class* = I, *Epitope Object Type* = Linear peptide). IEDB contains both quantitative and qualitative MHC-I epitope data, depending on whether the study used *in vitro* binding affinity assays or elution studies. Quantitative records were identified by *“Quantitative measurement”* !=“NA”, *dataset_type=”BA”* and *Units=”nM”*. Qualitative records were identified by *“Quantitative measurement”* =“NA” and *Units!=”nM”*. In order to add data that is not included in IEDB, we identified binding records released by MHCFlurry 2.0^16^ which were absent from IEDB. The MHCFlurry 2.0 datasets were downloaded from https://data.mendeley.com/datasets/zx3kjzc3yx/3. From the MHCFlurry mass spectrometry dataset, we first removed 52,642 entries corresponding to Class II MHC measurements. Next, records obtained via binding affinity assays, multi-allelic (MA) assays and single-allelic (SA) assays from both IEDB and MHCFlurry data were each filtered for overlap separately. MA records are peptide-MHC binding instances from experiments with cell lines naturally encoding multiple class-I MHC alleles, whereas SA records are derived from cell lines engineered to have a single class-I MHC allele. In the IEDB dataset, SA records are distinguished from MA records by *“Allele Evidence Code*”=*“Single allele present”*. For each data type (BA, SA and MA), MHCFlurry peptide-MHC records were added to the IEDB dataset if absent from the corresponding partition of the IEDB dataset. The MA dataset from MHCFlurry (S1 dataset) does not contain peptide-allele assignments, but rather best allele assignments by multiple pan-MHC binding tools. To determine best assignments to supplement our training dataset, we utilized MHC assignments which agree among all of three previously published methods, NetMHCpan 4.1 EL^11^, MHCFlurry 2.0 BA^16^ and MixMHCpred 2.0.2^14^. We ran NetMHCpan 4.1 EL to assign the best MHC to each peptide, and assignments for the other two methods were already published as columns within the S1 dataset. MHC allele assignments were in agreement among all three methods for 68,555 peptide-MHC observations out of the 91,581 observed epitopes in multiallelic data, and were added to our curated dataset after confirming the data was not in the IEDB MA dataset.

Further preprocessing steps were taken on the compiled epitope dataset. For quantitative records with multiple observations of a given peptide-MHC pair, the mean binding affinity value was assigned to the peptide-MHC pair. Records were excluded if the difference between maximum and minimum log-transformed values exceeded 0.2, as previously described^33^ [i.e., (1 − log(nM)/log(50,000)) > 0.2]. All positive qualitative records were assigned a quantitative affinity value of 100nM and measurement inequality of #x201C;<” as performed by O’Donnell *et. al*.^16^. The measurement inequalities (< and >) are used for semi-quantitative and qualitative records so that binding prediction models do not penalize predictions which are within the range of certainty for the associated binding affinity. All qualitative negatives were assigned a very low binding affinity value sampled between 20,000 and 50,000nM to increase diversity among samples during training, and assigned a measurement inequality of “>” to account for the uncertainty in the binding affinity value. Measurement inequalities “<=” and “>=” recorded in the IEDB were mapped to “<” and “>” respectively.

